# Transcranial ultrasound stimulation effect in the redundant and synergistic networks consistent across macaques

**DOI:** 10.1101/2023.11.02.564776

**Authors:** Marilyn Gatica, Cyril Atkinson-Clement, Pedro A. M. Mediano, Mohammad Alkhawashki, James Ross, Jèrôme Sallet, Marcus Kaiser

## Abstract

Low-intensity transcranial ultrasound stimulation (TUS) is a non-invasive technique that safely alters neural activity, reaching deep brain areas with good spatial accuracy. We investigated the effects of TUS at the level of macaque using a recent metric, the synergy minus redundancy rank gradient, that quantifies different kinds of causal neural information processing. We analyzed this high-order quantity on the fMRI data after TUS in two targets: the supplementary motor area (SMA-TUS) and the frontal polar cortex (FPC-TUS). The TUS produced specific changes at the limbic network at FPC-TUS and the motor network at SMA-TUS and altered, in both targets, the sensorimotor, temporal, and frontal networks, consistent across macaques. Moreover, there was a reduction in the structural and functional coupling after both stimulations. Finally, the TUS changed the intrinsic high-order network topology, decreasing the modular organization of the redundancy at SMA-TUS and increasing the synergistic integration at FPC-TUS.

## Introduction

Low-intensity transcranial ultrasound stimulation (TUS) is a promissory and non-invasive neuro-modulation technique that can safely alter neural activity and reach both cortical and deep areas with good spatial accuracy in comparison to other non-invasive brain stimulation methods (***Legon et al., 2020***; ***Bystritsky et al., 2011***; ***Darmani et al., 2022***). Although the exact mechanisms of TUS are still a matter of debate, some hypotheses have been suggested. At a microscopic level, TUS alters brain cells without causing a significant heating increase in the tissue (***Naor et al., 2016***) through mechanical stimulation of sodium and calcium channels (***Kubanek et al., 2016, 2018***; ***Tyler et al., 2008***) and/or microcavitation resulting in local depolarization and alteration of the glia neuron decoupling (***Krasovitski et al., 2011***; ***Plaksin et al., 2016***; ***Oh et al., 2019***). On a macroscopic level, TUS is related to an increased brain’s target temperature without causing oedema or impairing the blood-brain barrier (***Webb et al., 2023***) and to an increased excitability following a reduced GABA inhibition (***Yaakub et al., 2023***). TUS has also shown functional network alterations depending on the structural coupling of the target, with an increase in the brain functional connectivity with the strongly connected areas and a decrease in the correlations in the less connected regions (***Folloni et al., 2019***; ***Verhagen et al., 2019***) as well as behavioral (***Hameroff et al., 2013***; ***Fouragnan et al., 2019***; ***Sanguinetti et al., 2020***; ***Nakajima et al., 2022***; ***Mahmoodi et al., 2023***) changes from several minutes to several days following the stimulation.

However, studies are now required to understand how TUS could contribute to brain reorganization through high-order interdependencies explorations under the two types of interactions: redundancy and synergy. The redundancy can be understood as the information we can obtain in any system component. The synergy is the extra information we get if we only observe the whole system together. Several mathematical and computational tools have been proposed to estimate them, whereas different studies have pointed out their relevance (***Williams and Beer, 2010***; ***Timme et al., 2014***; ***Lizier, 2014***; ***Rosas et al., 2019***). A recent study reported the redundancy linked with lower-level sensorimotor processing and the synergy with higher-level cognitive tasks (***Luppi et al., 2022***). Furthermore, they exhibited different network organization, with the redundancy being more segregated and correlated with the structural connectivity. In contrast, the synergy was associated with the wiring distance matrix between pairs of regions and favored integrated processing (***Luppi et al., 2022***). In healthy aging, redundancy increased in the older population (***Camino-Pontes et al., 2018***; ***Gatica et al., 2021a***), and could be explained by a non-linear-neurodegenerative model applied to the structural connectivity (***Gatica et al., 2021b***). Furthermore, these high-order methods have been used in a wide range of studies, such as neurodegeneration (***Herzog et al., 2022***), artificial neural networks (***Proca et al., 2022***), spiking neurons (***Stramaglia et al., 2021***), and elementary cellular automata (***Rosas et al., 2018***; ***Orio et al., 2023***).

This article aims to elucidate how the TUS could reorganize the brain as measured by the computation of the causal redundancy and synergy at the individual level of macaques. In this direction, we re-analyze the TUS effect on resting-state fMRI data of three macaques under anesthesia, on a time period from 30 to 150 minutes following the stimulation (***Verhagen et al., 2019***). We independently characterized the high-order quantities at the control condition (non-TUS) and after applying TUS at two targets: the supplementary motor area (SMA-TUS) and the frontal polar cortex (FPC-TUS). Our results showed that the TUS produced target-specific changes in the rank gradient distribution at the limbic network at FPC-TUS and the motor network at SMA-TUS and alterations in common, independent of the target, on the sensorimotor, temporal, and frontal networks. Secondly, the differences were consistent across macaques. Thirdly, the TUS decreased the structural and functional coupling independent of the stimulated target. Finally, the TUS changed the intrinsic high-order network topology, reducing the modular organization of the redundancy at SMA-TUS and increasing the synergistic integration at FPC-TUS.

## Results

We first developed a simple example of two sinusoids to give an interpretation of how the redundancy and synergy change when modifying one of the signals. The sinusoids started as the same function, resulting in high redundancy and synergy zero (***Figure 1***A), and after modifying the second one gradually, the redundancy decreased, and the synergy increased (***Figure 1***B) until the interaction is purely synergistic (***Figure 1***C). In particular, we obtained high values of redundancy and synergy due to the noiseless system, and they would be smaller if some noise were added, although the intuitions would remain. However, both quantities were annulled when the second function was noise-dominated (***Figure 1***D). Therefore, if we modulated one of the signals, we observed that the redundancy and synergy nulled at low and complete synchrony, respectively. In contrast, there was a region where both quantities coexisted.

**Figure 1.**
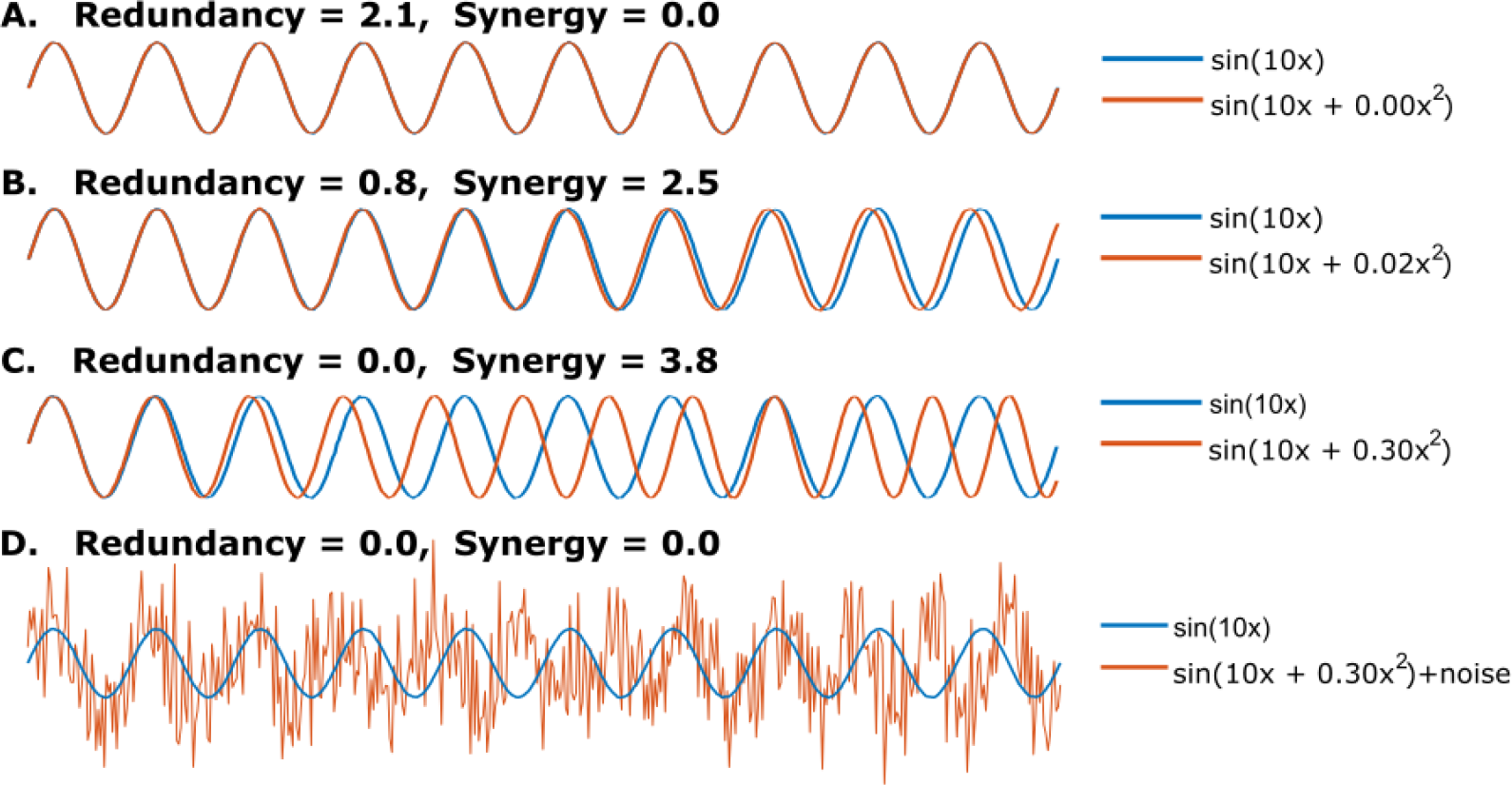
Given *f* (*x*) = *sin*(10*x*) and *g*(*x*) = *sin*(10*x* + *ix*^2^), with *i* = {0, 0.02, 0.3}, and *x ∈* [0, 2*π*]. **A**. full redundancy and zero synergy when *f* = *g*. **B**. The redundancy decreases when i = 0.02. **C**. Full synergy and zero redundancy when i = 0.3. **D**. The synergy and redundancy are zero when i = 0.3 and adding noise *∼* 𝒩 (0, 1.1).

Next, we analyzed fMRI data of three macaques at non-TUS, SMA-TUS, and FPC-TUS from 30 to 150 minutes following the stimulation (***Figure 2***A). To capture a region’s prevalence of redundancy or synergy, we performed a dynamical extension to the synergy minus redundancy rank gradient (***Figure 2***B-C), which considers causality while estimating high-order interactions (***Luppi et al., 2022***). Finally, to answer if some areas were shifted to more redundant or synergistic interactions after the stimulation, we compared the rank gradient distribution between each TUS experiment and the control condition per each brain region (***Figure 2***D).

**Figure 2.**
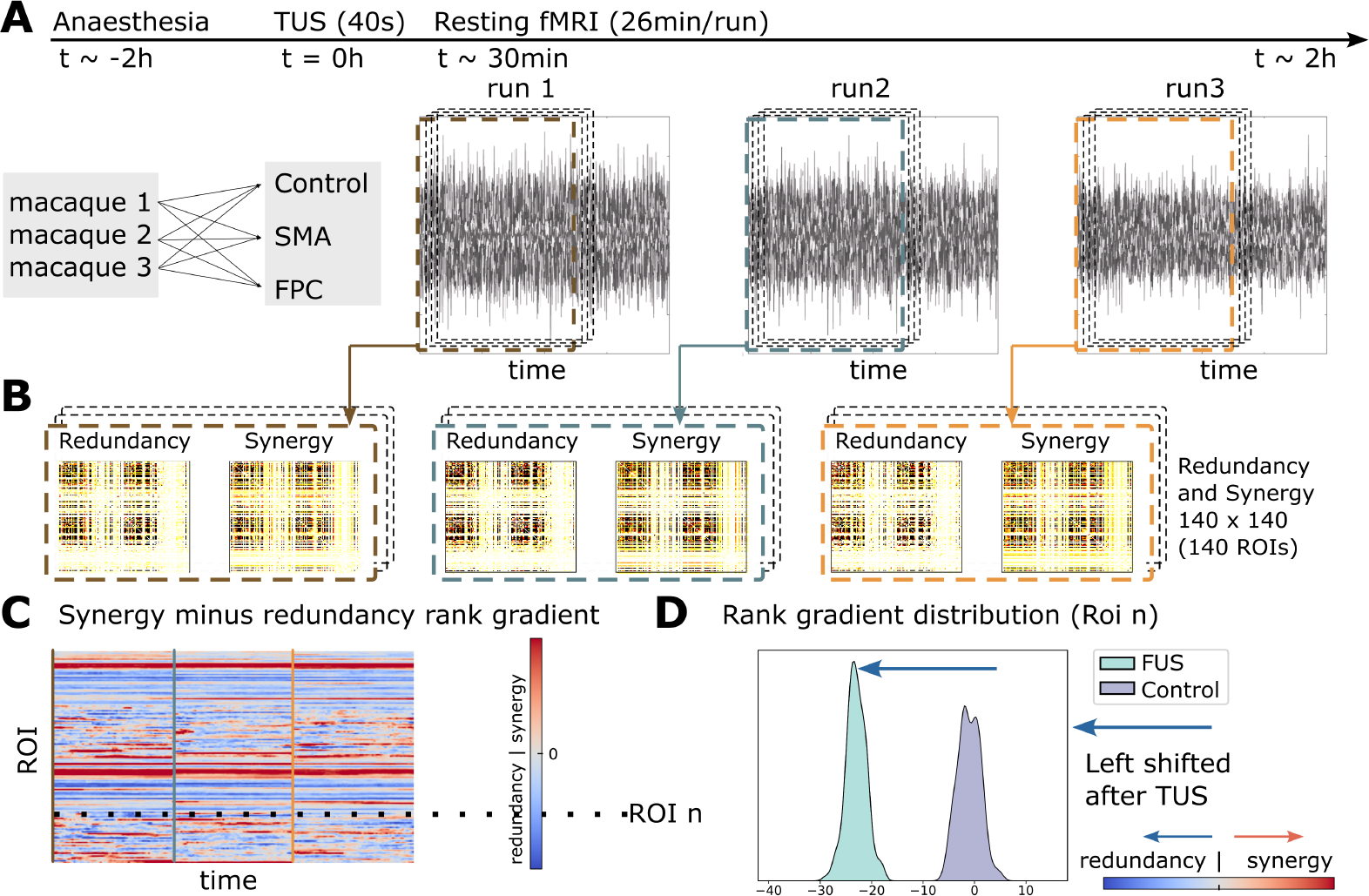
**A**. Three macaques participated in FPC-TUS, SMA-TUS, and non-TUS. **B**. We computed the redundancy and synergy matrices over 60 sliding windows of 500s. **C**. The matrices are averaged across rows and then ranked. The result of those synergy minus redundancy vectors is the rank gradient (***Luppi et al., 2022***). **D**. Per each ROI, we compared the gradient rank distribution over time (dotted line) between the control and TUS, obtaining a shift to more redundant (blue arrow) or synergistic (red arrow) interactions.

### Synergy-redundancy rank gradient disruption after SMA-TUS

To investigate the whole-brain effects of the stimulation at the SMA, we compared the synergy minus redundancy rank gradient between the non-TUS condition and the SMA-TUS. First, we grouped the three macaques and compared the gradient rank distribution between the control (non-TUS) and the SMA-TUS. Several networks were affected after the stimulation at SMA (***Figure 3***A). The sensorimotor area was altered in the right secondary somatosensory cortex (SII, es = −0.94) towards redundancy and the left inferior parietal lobule (area_7_in_IPL, es = 0.92) to synergy. The temporal area in the left fundus of the superior temporal sulcus (STSf, es = −0.80) participated in more redundant interactions. The frontal cortex, particularly the right lateral orbital frontal cortex (lat_OFC, es = 1.03) and the right ventrolateral prefrontal cortex (vlPFC, es = 1.80) shifted towards synergy. The motor area was altered to redundancy in the right pallidum (Pd, es = −0.87) and left posterior thalamus (PThal, es = −1.06). The effect size comparisons (SMA-TUS minus non-TUS) of those regions are shown on brain maps (***Figure 3***B). Moreover, at the individual level, the differences are homogeneous across two or three macaques (***Figure 3***C). Therefore, we found differences between non-TUS and SMA-TUS across the sensorimotor (↑ redundancy and ↑ synergy), frontal (↑ synergy), temporal (↑ redundancy), and motor (↑ redundancy) networks at group and individual levels.

**Figure 3.**
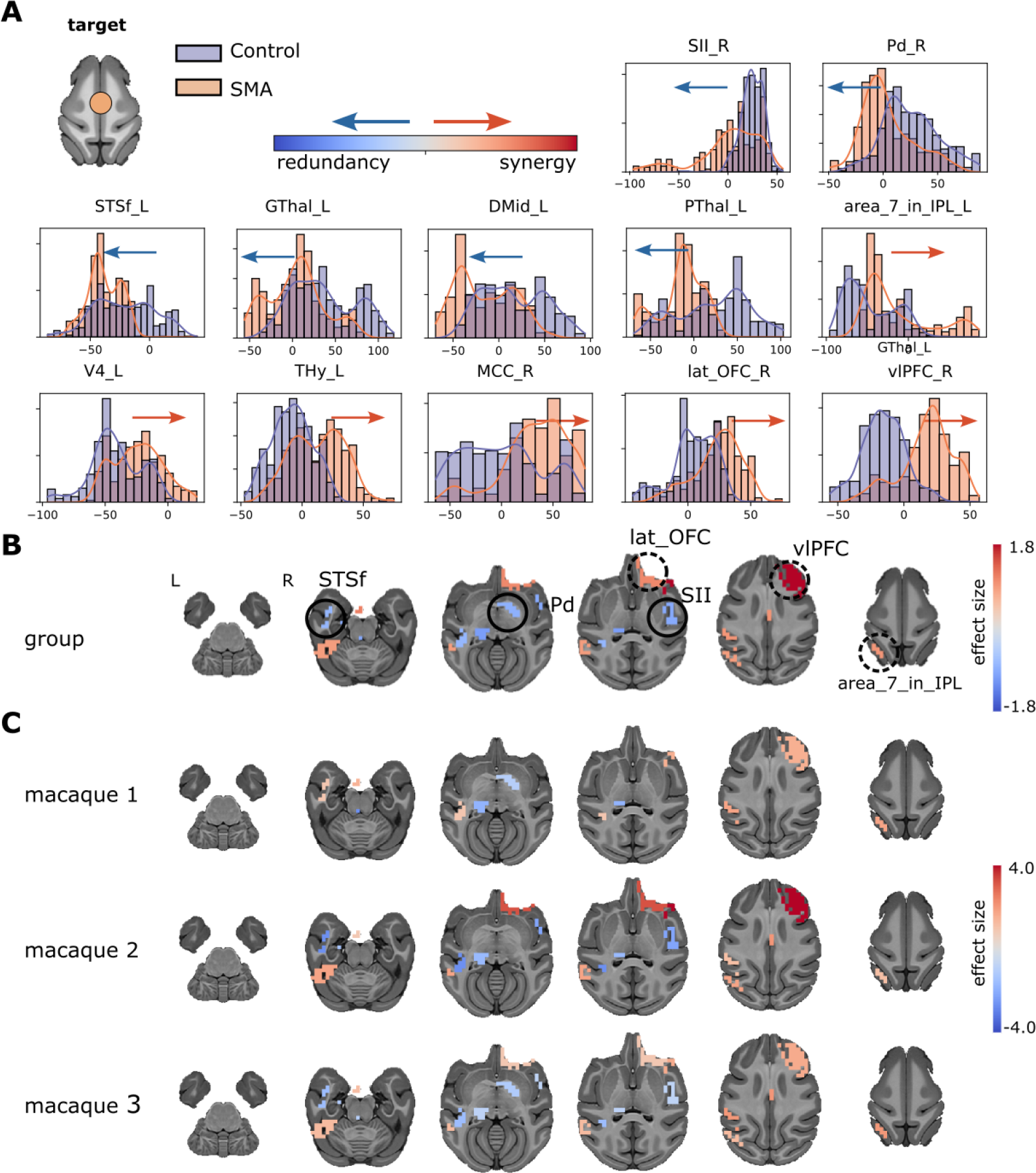
**A**. Synergy minus redundancy rank gradient distribution after the TUS of the SMA target. The left shift (blue arrow) represents a region participating in more redundant (or less synergistic) interactions over time after TUS. In contrast, the red arrow (right shifting) represents a more synergistic (or less redundant) interaction over time after TUS. **B**. The brain maps illustrate the shift to the left (blue color) or right (red color), and the magnitude is the effect size. We compared the gradient rank distribution of each ROI over time between the control (non-TUS) and the TUS experiment among the three macaques together, using a Wilcoxon rank-sum test and correcting by Bonferroni and effect size bigger than 0.8. **C**. Similar to B, but for each macaque separately.

### Synergy-redundancy rank gradient disruption after FPC-TUS

To explore the network alterations of the stimulation at FPC, we compared the synergy minus redundancy rank gradient between the control condition and the FPC-TUS. We found differences in the gradient rank distribution across several networks at the group level (***Figure 4***A). The sensorimotor network incremented the redundancy at the right secondary somatosensory cortex (SII, es = −1.07). In contrast, the sensorimotor region was altered towards synergy at the right primary somatosensory cortex (SI, es = 1.21) and the left inferior parietal lobule (area_7_in_IPL, es = 1.02). The temporal area shifted to redundancy at the left caudal superior temporal gyrus (STGc, es = −0.91). The frontal cortex switched towards synergy at the right lateral orbital frontal cortex (lat_OFC, es = 0.87). The limbic network shifted to redundancy at the left striatum (Str, es = −0.99), right hippocampus (paraHipp, es = −1.23), and right anterior cingulate cortex (ACC, es = −0.89). Additionally, The limbic network changed towards synergy at the left ventral midbrain (VMid, es = 1.03). The effect size comparisons of those regions are shown on brain maps (***Figure 4***B) for the group analysis. Most differences are consistent across two or three macaques, except the left amygdala, left pons, right lateral orbital frontal cortex, and right hypothalamus, which were statistically significantly different at only one macaque (***Figure 4***C). In conclusion, the stimulation at the FPC produced an effect on the somatosensory (↑ redundancy and ↑ synergy), temporal (↑ redundancy), frontal (↑ synergy), and limbic (↑ redundancy and ↑ synergy) networks.

**Figure 4.**
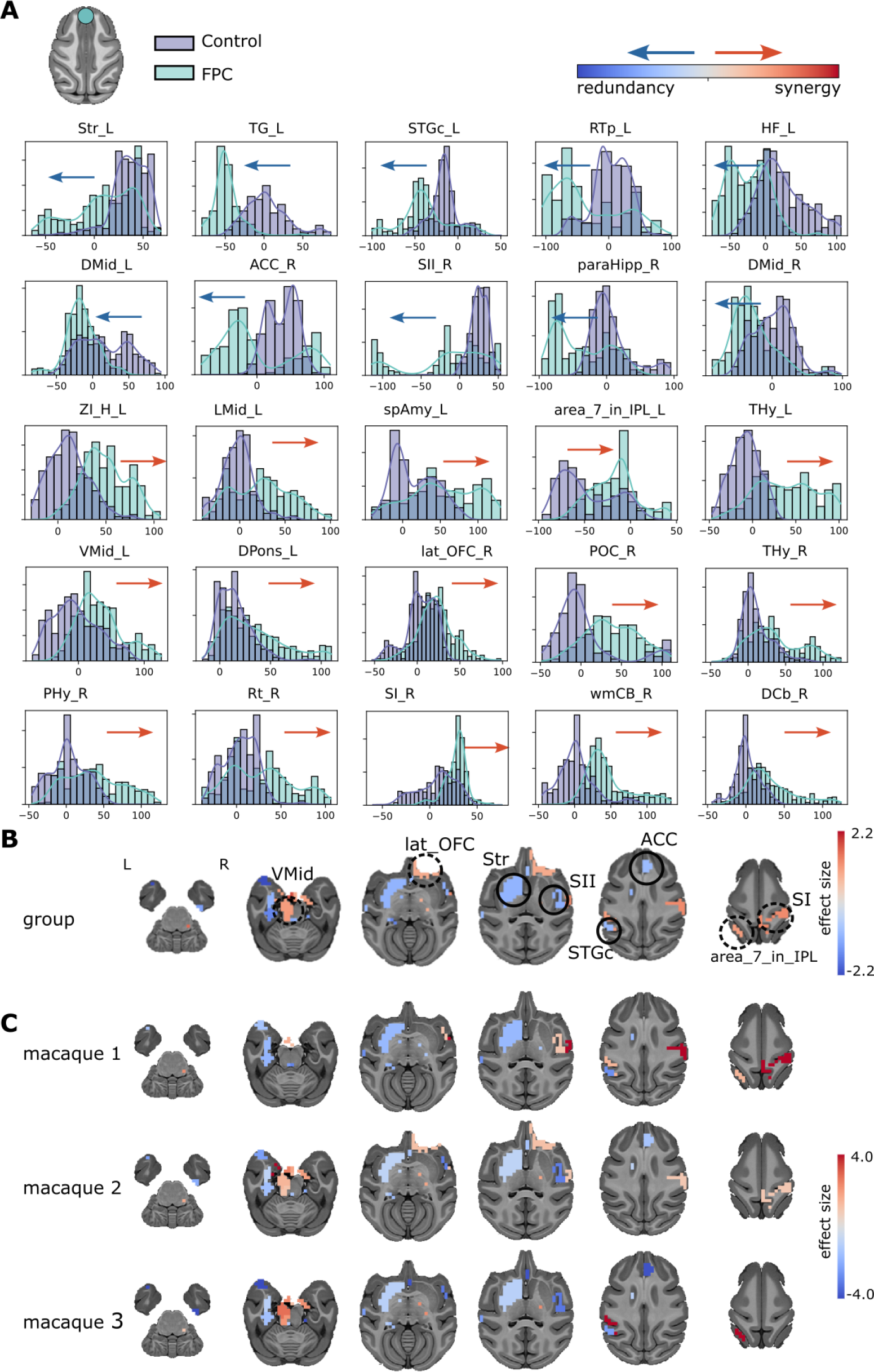
**A**. Synergy minus redundancy rank gradient distribution after the TUS of the FPC target. The left shift (blue arrow) represents a region participating in more redundant (or less synergistic) interactions over time after TUS. In contrast, the red arrow (right shifting) represents a more synergistic (or less redundant) interaction over time after TUS. **B**. The brain maps illustrate the shift to the left (blue color) or right (red color), and the magnitude is the effect size. We compared the gradient rank distribution of each ROI over time between the control (non-TUS) and the TUS experiment among the three macaques together, using a Wilcoxon rank-sum test and correcting by Bonferroni and effect size bigger than 0.8. **C**. Similar to B, but for each macaque separately.

### Structural and functional coupling

To understand if there is an alteration in the functional and structural coupling produced by TUS, we quantified the similarities between the high-order quantities with the structural connectivity (SC) and the Euclidean distance (ED). The SC-redundancy correlation in controls was *ρ* =0.24 and decreased to *ρ* =0.198 at FPC-TUS. Moreover, it decayed to *ρ* =0.196 at SMA-TUS, on group average (***Figure 5***A), and those differences persisted in all the macaques. Likewise, the ED-synergy correlation at non-TUS was *ρ* = 0.20 and decreased to *ρ* = 0.11 after FPC-TUS. Additionally, the differences were consistent in the macaques two and three (***Figure 5***B). Nevertheless, the similarity increased from *ρ* = 0.20 (non-TUS) to *ρ* = 0.24 at SMA-TUS, on average, across macaques. The consistent differences were observed in the macaques one and three (***Figure 5***B). Finally, the ED-synergy correlation was more significant than SC-synergy, and there was no apparent disruption in SC-synergy after TUS (***Figure 5—figure Supplement 1***A). Similarly, the SC-redundancy correlation was more prominent than ED-redundancy and without significant disruptions at the ED-redundancy after any stimulation (***Figure 5—figure Supplement 1***B). In conclusion, the SC-redundancy correlation decreased independent of the target. The ED-synergy similarity increased at SMA-TUS and decayed at FPC-TUS.

**Figure 5.**
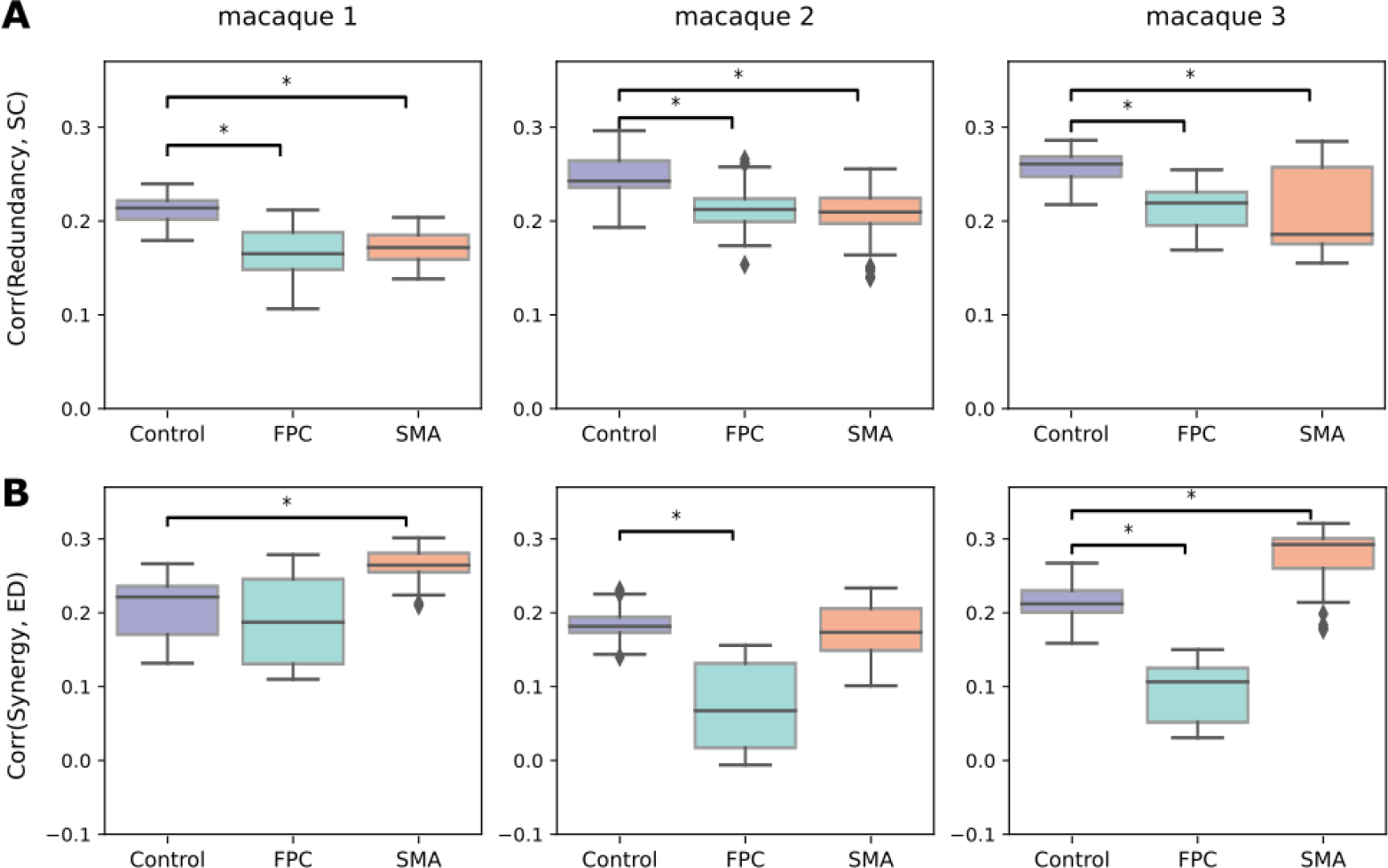
**A:** Correlation between the structural connectivity (SC) and the redundancy. **B:** Correlation between the Euclidean distance (ED) and synergy per experiment and macaque (at each column). The y-axis values contain the Spearman’s rank correlation coefficient. The colors represent the control (non-TUS) and the two experiments: SMA-TUS and FPC-TUS. **Figure 5—figure supplement 1**. **A:** Correlation between the structural connectivity (SC) and the synergy. **B:** Correlation between the Euclidean distance (ED) and redundancy per experiment and macaque.

### Network reorganization after TUS

To further understand if a network reorganization exists in the functional high-order interactions, we seek to characterize the network topology of the redundancy and synergy before and after the stimulation. We used modularity and global efficiency based on the pioneer topological characterization of redundancy and synergy (***Luppi et al., 2022***). The redundancy exhibited a modular organization (***Figure 6***A), and the synergy presented an integrated topology (***Figure 6***B). At the same time, the modularity in the synergy and the global efficiency in the redundancy were near zero in non-TUS or TUS experiments. Interestingly, the modularity of the redundancy decreased after the SMA stimulation (***Figure 6***A) in two macaques, with unclear changes in the synergistic global efficiency (***Figure 6***B). In contrast, the FPC-TUS increased the synergistic integration (***Figure 6***B) without consistent alteration at the modular organization in the redundancy (***Figure 6***A). Altogether, the intrinsic modularity in the redundancy decreased after SMA-TUS, and the segregation in the synergy increased after FPC-TUS.

**Figure 6.**
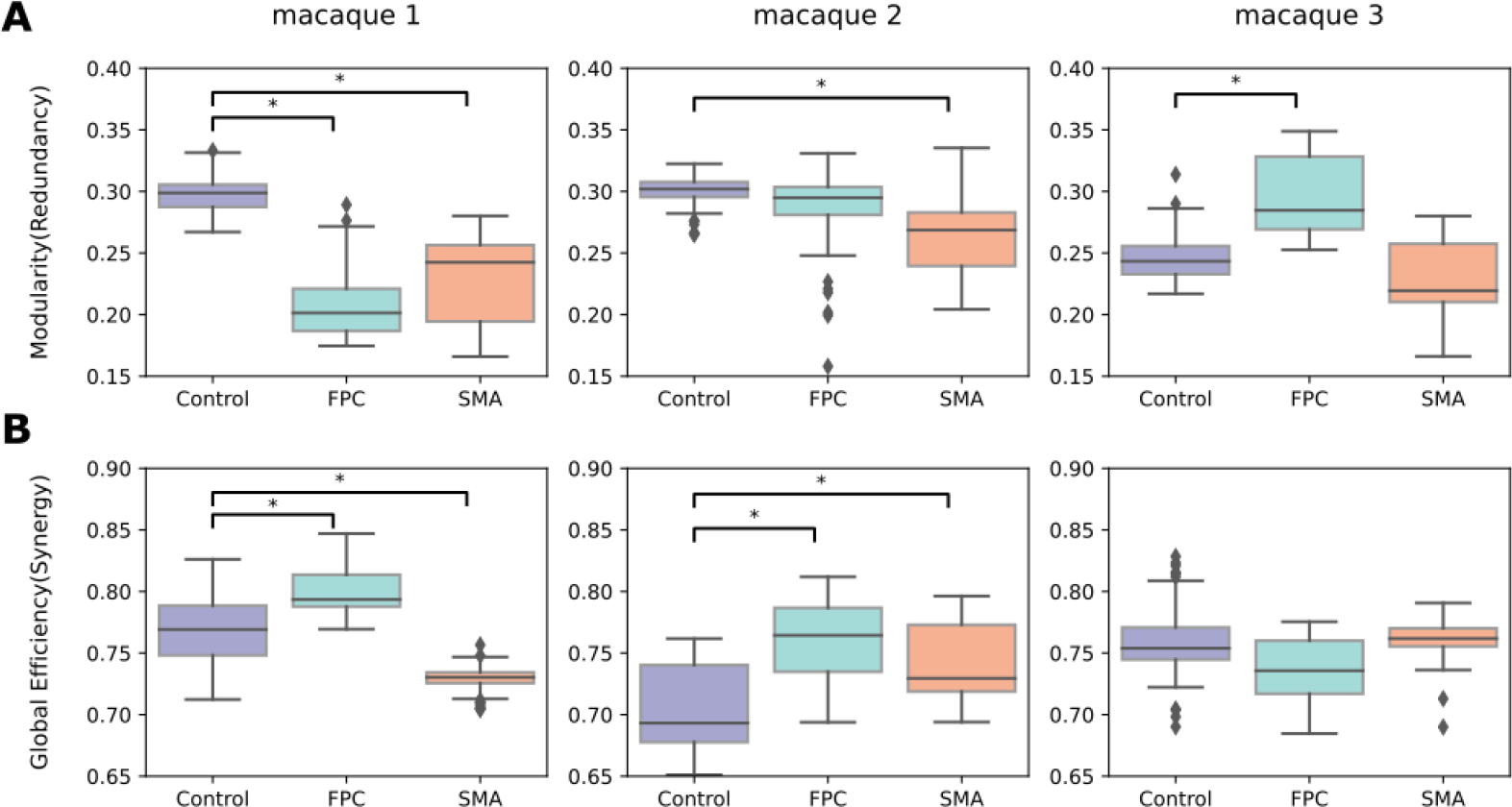
**A**: Modularity (segregation) of the redundancy matrix. **B:** Global efficiency (integration) of the synergy matrix. The colors represent the experiment or control condition, and each column depicts a macaque.

## Discussion

This article presented three main results: 1. The TUS produced high-order changes depending on the target at the limbic network at FPC-TUS and the motor network at SMA-TUS and altered in both targets, the sensorimotor, frontal, and temporal networks (***Figure 7***A). 2. The differences were consistent across macaques. 3. The TUS decreased the functional and structural coupling independent of the targets and modified the intrinsic high-order topological organization. SMA-TUS decreased the modularity of the redundancy, and the FPC-TUS increased the synergistic integration (***Figure 7***B).

**Figure 7.**
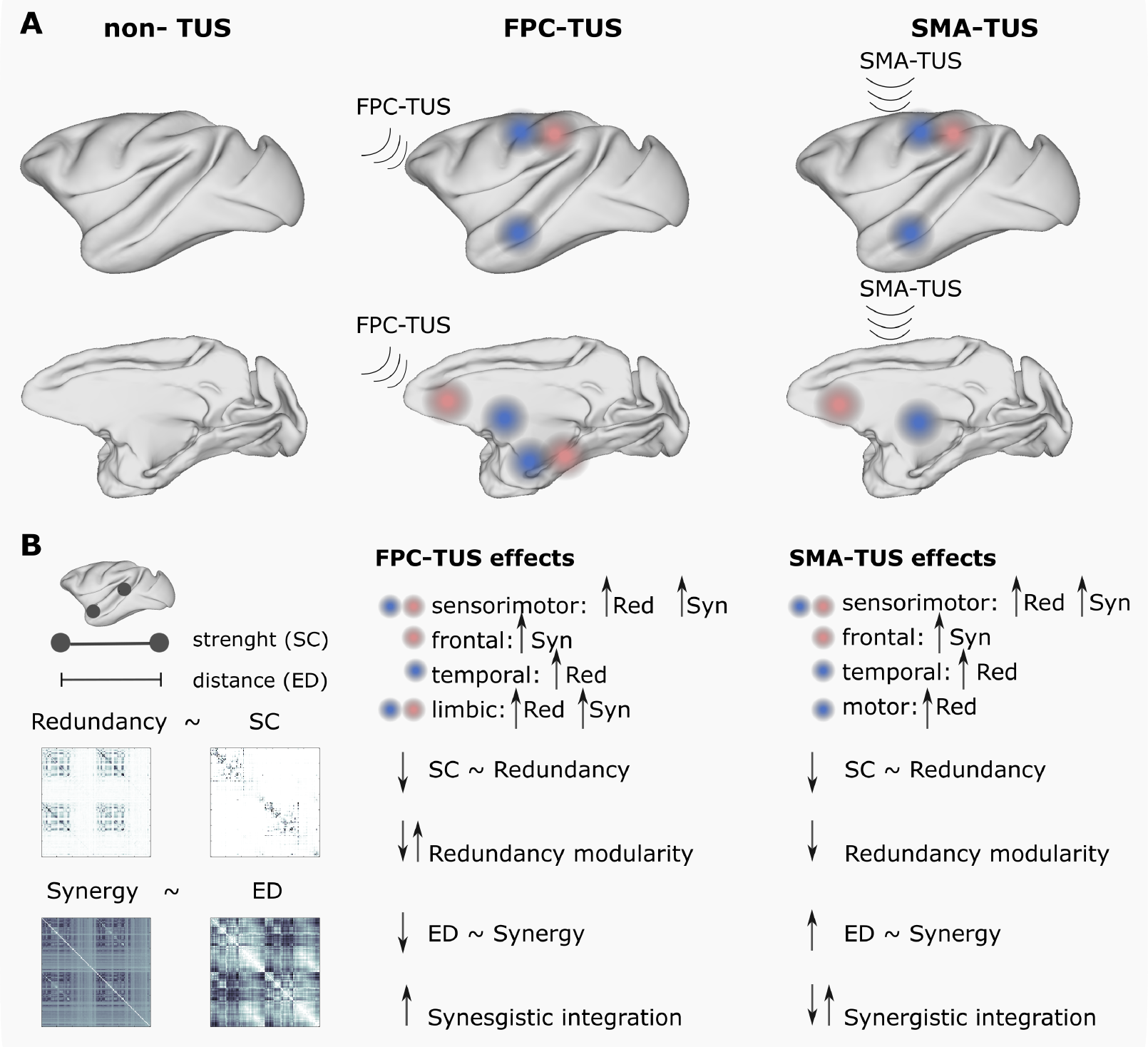
Overview. **A** The TUS produced target-specific changes in the rank gradient distribution at the limbic network at FPC-TUS and the motor network at SMA-TUS and alterations in common, independent of the target, on the sensorimotor, temporal, and frontal networks. **B:** The TUS decreased the structural and functional coupling and altered, depending on the target, the intrinsic high-order network topology.

### SMA-TUS changes the rank gradient consistently across macaques

We found some regions changing to more redundant interaction at the sensorimotor (secondary somatosensory cortex), temporal (fundus of the superior temporal sulcus), and motor networks (pallidum and thalamus). In contrast, some networks presented a shift to synergistic interactions, such as the sensorimotor (inferior parietal lobule) and frontal (ventrolateral prefrontal cortex and lateral orbital frontal cortex) networks (***Figure 3***). Although we used a causal high-order analysis instead of Pearson correlations, some regions are consistent with the previous study that analyzed the same data, reporting changes in the sensorimotor networks, prefrontal cortex, anterior temporal, and anterior and posterior cingulate at SMA-TUS on macaques. (***Verhagen et al., 2019***). Moreover, targeting motor circuits on transcranial magnetic stimulation (TMS) produces changes not only in the somatosensory network (***Bestmann et al., 2004***; ***Jung et al., 2020***) but also spread across different areas, for example, in the lateral frontotemporal cortex, including the inferior frontal gyrus (***Pineda-Pardo et al., 2019***), on human fMRI data.

### FPC-TUS changes the rank gradient consistently across macaques

There are some networks shifted to more redundant interaction at FPC-TUS, such as the sensorimotor (secondary somatosensory cortex), temporal (the caudal superior temporal gyrus), and limbic (striatum, hippocampus, and anterior cingulate cortex). In contrast, other networks switched to more synergistic interdependencies, such as the sensorimotor (primary somatosensory cortex and the inferior parietal lobule), frontal (lateral orbital frontal cortex), and limbic (ventral midbrain). (***Figure 4***). Even though we assessed a causal analysis instead of Pearson correlations, several of these areas corresponded with the previous study that analyzed the same macaque fMRI data showing changes in the lateral prefrontal areas, the superior temporal sulcus, the posterior cingulate cortex, and the sensorimotor networks presented functional connectivity differences after the FPC-TUS (***Verhagen et al., 2019***). On the other hand, the limbic network, particularly the striatum, thalamus, and amygdala, has modulated their functional connectivity after TMS at the frontopolar cortex in human data (***Riedel et al., 2019***; ***Hanlon et al., 2013***). That network is also relevant for macaques, where the prefrontal cortex and the limbic network are widely connected (***Carmichael and Price, 1995***; ***Petrides and Pandya, 2007***).

### TUS decreases the structure-function coupling

The redundancy showed a stronger correlation with the SC, whereas the synergy with the distance consistently with previous studies (***Luppi et al., 2022***). Those similarities were disrupted after TUS, with a global decrease in the correlations, except for the distance and synergy correlation that increased after SMA-TUS (***Figure 5***). The findings indicated that there may have been a high-order network reorganization driven by stimulation. At the level of pairwise correlations, previous studies have shown changes in the functional connectivity after TUS depending on the structural coupling of the target (***Verhagen et al., 2019***; ***Folloni et al., 2019***; ***Bestmann et al., 2004***; ***Pineda-Pardo et al., 2019***), with an increase in the functional connectivity in the strongly connected areas, which are usually near to the target.

### TUS altered the high-order intrinsic organization

The redundancy presented a modular organization, whereas the synergy showed an integrated topology (***Figure 6***), consistent with previous studies (***Luppi et al., 2022***; ***Varley et al., 2023***). A seminal study described the integration and segregation as two underlying processes of brain organization that co-exist, allowing to perform diverse cognitive tasks (***Tononi et al., 1994***). Lower cognitive tasks have been linked with higher functional segregation in simple motor tasks. However, in working memory tasks have been reported an increase of integration (***Cohen and D’Esposito, 2016***), where the prefrontal cortex had a critical role (***Diamond, 2013***; ***Menon and D’Esposito, 2022***). In terms of high-order, lower-level processing networks such as the motor area had been liked with a prevalence of redundancy, whereas the frontal cortex was predominated by synergy (***Luppi et al., 2022***). Interestingly, our findings also showed different high-order reorganizations depending on whether the target is the supplementary motor area or the prefrontal cortex, with the TUS altering the modular organization of the redundancy at SMA-TUS and, in contrast, increasing the intrinsic integration in the synergy at FPC-TUS.

### Limitations and future work

The current study has some limitations. First, rather than using absolute values, the synergy minus rank gradient depends on relative values dominated by redundancy and synergy. Secondly, only three macaques were examined in this study, and larger sample sizes should be used in further research. Third, we analyzed macaque fMRI data under anesthesia. Previous research has demonstrated that each anesthesia treatment altered functional connectivity differently in the rat (***Paasonen et al., 2018***), macaque (***Giacometti et al., 2022***), and human brain (***Peltier et al., 2005***). Nevertheless, macaques were anesthetized using inhalational isoflurane gas based on a widely used protocol that preserves whole-brain functional connectivity (***Vincent et al., 2007***; ***Sallet et al., 2013***; ***Neubert et al., 2015***). Finally, resting-state fMRI data for human participants should be evaluated in future studies to move toward employing neuromodulation for therapeutic purposes.

## Conclusion

To the best of our knowledge, it is the first high-order analysis of causal interactions after TUS. Our results indicate that the causal interactions are altered after TUS with consistencies across macaques, and the patterns of changes depend on the target. Although using high-order interactions needs further research in TUS, they have been explored in many applications, and might also be a relevant methodology in TUS experiments as a complementary approach to Pearson correlation.

## Methods and Materials

### Ultrasound stimulation

The ultrasound transducer device was a 64 mm diameter, H115-MR, Sonic Concept, Bothell, WA, USA, with 51.74 mm focal depth, used with a coupling cone sealed with a latex membrane (Durex) and filled with degassed water. The protocol was controlled with a digital function generator (Handyscope HS5, TiePie engineering, Sneek, Netherlands) setting to 250 kHz with 30 ms bursts of ultrasound generated every 100 ms. A 75-watt amplifier (75A250A, amplifier research, Souderton) was used to deliver the power to the transducer. A TiePie probe (Handyscope HS5, TiePie engineering, Sneek, The Netherlands), connected to an oscilloscope, was used to monitor the voltage. The recorded peak-to-peak voltage remained constant throughout the stimulation. Per session, the voltages ranged from 130-142 V, analogous to 1.17-1.35 MPa, as measured in water with an in-house heterodyne interferometer (***Constans et al., 2017***). For the SMA target, the maximum peak pressure (P max) and I sspa at the acoustic focus point estimations were 0.88MPa and 24.1 W/cm 2 (I spta : 7.2 W/cm 2). For the FPC target, the parameters were 31.7 W/cm 2(I spta : 9.5 W/cm 2) (***Verhagen et al., 2019***). The stimulation lasted for 40 seconds. The two target locations were close to the midline, stimulating both hemispheres simultaneously with a single train. The stimulation was guided using a frameless stereotaxic neuronavigation system (Rogue Research, Montreal), registering a T1-weighted MR image to each macaque’s head. The (X, Y, Z) Montreal Neurological Institute (MNI) coordinates were (0.1, 2,19) at SMA and (−0.7, 24, 11) at FPC. The positions of the transducer and the animal head were tracked continuously using infrared reflectors. The transducer was placed on previously shaved skin using conductive gel (SignaGel Electrode; Parker Laboratories Inc.) to ensure ultrasonic coupling between the transducer and the scalp. There were at least ten days in between two TUS sessions. In the control condition (non-TUS), all procedures, including anesthesia, pre-scan preparation, fMRI scan acquisition, and timing, with the exception of actual TUS, matched with the TUS sessions.

### Macaque data acquisition

We used the offline TUS fMRI dataset of macaques from the repository https://git.fmrib.ox.ac.uk/lverhagen/offlinetus. The data consists of three MRI sessions per each Control (non-TUS), SMA-TUS, and FPC-TUS, and macaque (N=3). Each macaque was anesthetized using inhalational isoflurane gas (***Vincent et al., 2007***; ***Sallet et al., 2013***; ***Neubert et al., 2015***). Moreover, the macaques received injection of ketamine (10 mg/kg, intramuscular), xylazine (0.125–0.25 mg/kg, intramuscular), midazolam (0.1 mg/kg, intramuscular), atropine (0.05 mg/kg, intramuscularly), meloxicam (0.2 mg/kg, intravenously), and ranitidine (0.05 mg/kg, intravenously). Following the stimulation, the animals were placed in a sphinx position in a 3T MRI scanner (four-channel phased-array coil, Dr. H. Kolster, Windmiller Kolster Scientific, Fresno, USA). To avoid ketamine’s clinical peak, scanning started about two hours after anaesthesia. Intermittent positive pressure ventilation was kept up to ensure a steady respiration rate. VitalMonitor software (Vetronic Services Ltd.) was used to record and monitor the values of respiration rate, inspired and expired CO2, and isoflurane concentration. Additionally, the core temperature and SpO2 were monitored constantly.

The fMRI data were acquired for three runs of 26 min each approximately using the scans parameters: 36 axial slices; in-plane resolution = 2 × 2 mm, slice thickness = 2 mm, without slice gap, TR = 2000 ms, TE = 19 ms, and 800 volumes per run. A T1-weighted structural MRI was scaned per each macaque. Image acquisition was performed following a T1 weighted magnetization-prepared rapid-acquisition gradient echo sequence with voxel resolution = 0.5 × 0.5 × 0.5 mm.

Finally, a T1-weighted structural MRI scan was acquired using a T1-weighted magnetization-prepared rapid-acquisition gradient echo sequence (voxel resolution:0.5×0.5×0.5 mm), a Black bone (voxel resolution: 0.5×0.5×0.5 mm), and a DWI (voxel resolution: 1×1×1 mm) scans, per macaque.

### fMRI pre-processing

The fMRI preprocessing was performed following previous pipelines using AFNI (***RW., 1996***; ***RW and JS., 1997***): first, the T1 structural image was realigned to match the resting state. Next, the skull of the T1 image was removed. The cerebrospinal fluid (CSF) and grey and white matter were segmented and realigned to the template space. Next, a slice-time correction, despike, and motion correction were applied to the fMRI data, and each volume was aligned with the mean volume, and a motion correction was applied. Then, the preprocessed fMRI data was spatially normalized to the NMT v2.0 brain template, and the CHARM (***Reveley et al., 2017***; ***Jung et al., 2021***) and SARM (***Hartig et al., 2021***) atlases (level 3) were applied. Next, we detrended the fMRI using motion as a nuisance variable. Therefore, the original time series were grouped into 140 brain regions, and finally, a band-pass filter between 0.0025 and 0.05 Hz was applied (***Barttfelda et al., 2015***).

### Diffusion pre-processing

The diffusion data were preprocessed using MRtrix. The pipeline consisted of denoising, Gibbs ringing artifacts removing, Eddy current, and bias field correction. Next, probabilistic tractography was performed using multi-shell multi-tissue Constrained Spherical Deconvolution (CSD). First, 10 Million streamlines were generated with the grey matter-white matter interface optimization. Then, we applied a spherical-deconvolution-informed filtering of tractograms and reduced the number of streamlines to 1 Million. Next, the connectome was computed on the CHARM (***Reveley et al., 2017***; ***Jung et al., 2021***) and SARM (***Hartig et al., 2021***) atlases (level 3), resulting in a 140 × 140 structural connectivity (SC) matrix, each per macaque, by counting the number of white matter streamlines connecting all module pairs.

### Synergy minus redundancy rank gradient

Partial Information Decomposition (PID) Given three random variables, two source variables *X*^*i*^ and *X*^*j*^, and a target *Y*, the Partial Information Decomposition (***Williams and Beer, 2010***) is given by:

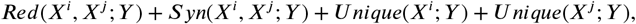

being *Red*(*X*^*i*^, *X*^*j*^ ; *Y*) the information provided by *X*^*i*^ and *X*^*j*^ about *Y* (redundancy), *Syn*(*X*^*i*^, *X*^*j*^ ; *Y*) is the information provided by *X*^*i*^ and *X*^*j*^ together about *Y* (synergy), *U nique*(*X*^*i*^; *Y*) is the information that is provided only by *X*^*i*^ about *Y*, and *U nique*(*X*^*j*^ ; *Y*) is the information that is provided only by *X*^*j*^ about *Y*. The PID could be represented by a forward decomposition into the nodes: 𝒜 = {{12}, {1}, {2}, {1}{2}}, being the nodes the synergistic, unique in source one, unique in source two, and redundant information, respectively.

#### Integrated information decomposition (Φ*ID*)

Consider the stochastic process of two random variables 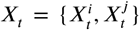 and denote the two variables in a current state *t*, by 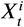 and 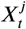, and the same two variables in a past state *t* − τ, by 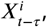, and, 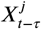. The integrated information decomposition (Φ*ID*) is the forward and backward decomposition of 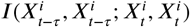, called the time delay mutual information, in redundant, synergistic and unique information (***Mediano et al., 2021***). Therefore, the Φ*ID* could be represented with the forward and backward interactions of the product 𝒜 *x* 𝒜. It results in 16 atoms (synergy to synergy, redundancy to redundancy, unique in source one to unique in source two (and backward), and redundancy to synergy, to name a few). In this article, we are focused on two atoms: persistent redundancy (redundancy that continues being redundancy) and persistent synergy (synergy that continues being synergy).

#### Synergy minus redundancy rank gradient

We performed a dynamical extension to the synergy minus redundancy rank gradient (***Luppi et al., 2022***). First, we computed the redundancy and synergy matrices measured with integrated information decomposition (Φ*ID*) over each sliding window. The Φ*ID* were computed over all the pairwise BOLD signal combinations 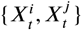, being *i* and *j* two different brain regions, with (*i, j*) *∈* {1, …, 140}, with the time series truncated to each sliding window. Next, each redundancy and synergy matrix belonging to the same window was averaged across rows separately, obtaining two strength vectors (each 1×140) and ranking their participation based on their strength. The rank gradient consists of the synergy minus redundancy rank, obtaining one vector (1×140) per window. In particular, each run lasted around 26 minutes, with 800 volumes or time points. We defined 500-time points per window with 99% overlap, resulting in 60 windows. As every session (TUS or non-TUS) and each macaque included three runs, we concatenated the synergy minus redundancy rank gradient of all the 3×60 = 180 windows, resulting in a matrix (140×180) per macaque and experiment. We compared the three macaque sessions concatenated (dim 140×540 each gradient rank matrix) for the global analysis at non-TUS vs. each TUS experiment. In contrast, for the case of individual comparisons, we used only the gradient rank of each macaque control vs. the TUS experiment (dim 140×180 each).

### Similarity

This analysis followed previous findings reporting the different structural support for the high-order quantities, where the redundancy was correlated with the structural connectivity (SC) and the synergy with the distance (***Luppi et al., 2022***). Because the SC is a sparse matrix, the structurefunction correlations were computed over the connected nodes. Therefore, the redundancy and synergy matrices were thresholded over the non-zero weight of the SC. Then, Spearman’s rank correlation coefficient was assessed between the upper triangular part of the thresholded SC and the upper triangular part of the redundancy (or synergy) matrix. In contrast, no thresholds were applied when comparing the distance and high-order correlations. We computed Spearman’s rank correlation coefficient between the upper triangular parts of the Euclidean distance matrix and the redundancy (or synergy).

### Network analysis

We performed the graph analysis for weighted networks using the Brain Connectivity Toolbox implementations (***Rubinov and Sporns, 2010***) over the redundancy and synergy matrices that, by definition, have non-negative values.

#### Segregation

To quantify segregation, we used the *modularity*, which enables the subdivision of the network into non-overlapping modules densely interconnected within each cluster and weakly connected between modules. The *modularity (Q)* were estimated using Newman’s spectral community detection algorithm (***Newman, 2006***; ***Reichardt and Bornholdt, 2006***). Mathematically, for a weighted graph, it is defined as:

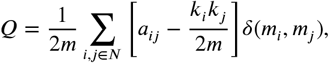

where *k*_*i*_ is the degree of the node *i, m*_*i*_ is the community of the node *i, m* is the sum of all of the edges in the graph, and *δ* is the Kronecker delta function (*δ* (*x, y*) = 1 if *x* = *y*, 0 otherwise).

#### Integration

To characterize the integration, we quantified the *global efficiency*, which is the inverse of the average shortest path length connecting two regions (***Latora and Marchiori, 2001***), meaning that, for disconnected nodes, their efficiency is zero. The *global efficiency* (*E*) is defined as:

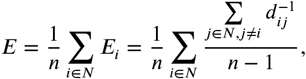

with *E*_*i*_ the efficiency of node *i*, and *d*_*ij*_ is the shortest path connecting the node *i* with the node *j*.

### Statistical analyses

This study compared the control (non-TUS) versus each TUS experiment (FPC-TUS or SMA-TUS). First, we performed a global analysis between the control and each TUS experiment, grouping the three macaques together and then at the level of macaque. The non-parametric statistical Wilcoxon rank sum test assessed the group and individual differences. We used Bonferroni correction and considered only the differences with an effect size bigger than 0.8. Finally, for the macaque-level analysis, besides Bonferroni and effect size correction, we constrain the areas showing differences to the regions belonging to the group mask of differences.

## Code availability

The data analysis was performed in MATLAB version 2022b. The MATLAB code to quantify synergy and redundancy from integrated information decomposition of time series with the Gaussian MMI solver is available at https://doi.org/10.1038/s41593-022-01070-0 (***Luppi et al., 2022***). The MATLAB code to assess the network analysis is freely available at brain-connectivity-toolbox.net (***Rubinov and Sporns, 2010***). We download the 3D macaque brain template from The Scalable Brain Atlas web page http://scalablebrainatlas.incf.org/ (***Bakker et al., 2015***; ***Markov et al., 2014***).

## Acknowledgments

M.K., J.R., and C.A. were supported by the Engineering and Physical Sciences Research Council (EP/W004488/1 and EP/X01925X/1). M.K. was also supported by the Guangci Professorship Program of Rui Jin Hospital (Shanghai Jiao Tong University). J.S. is funded by a Sir Henry Dale Wellcome Trust Fellowship 105651/Z/14/Z, IDEXLYON IMPULSION 2020 grant (IDEX/IMP/2020/14) and French National Research Agency grant (ANR-22-CE37-0021). The Wellcome Centre for Integrative Neuroimaging is supported Wellcome Trust core funding 203139/Z/16/Z.

---

**Figure S5.—figure supplement 1. A:**
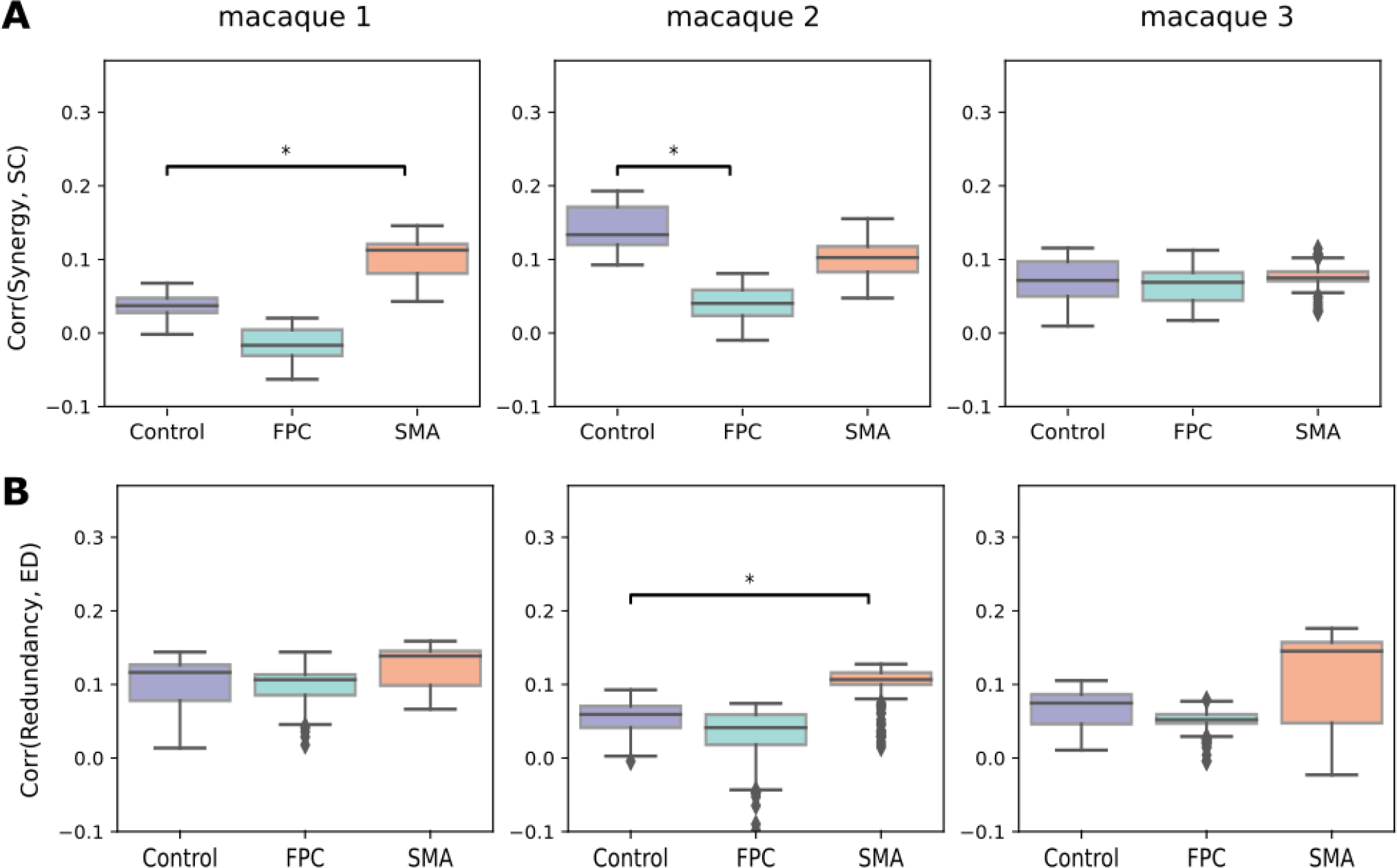
Correlation between the structural connectivity (SC) and the synergy. **B:** Correlation between the Euclidean distance (ED) and redundancy per experiment and macaque.

